# Discovery of ciliary G protein-coupled receptors regulating pancreatic islet insulin and glucagon secretion

**DOI:** 10.1101/2020.10.21.349423

**Authors:** Chien-Ting Wu, Keren I. Hilgendorf, Romina J. Bevacqua, Yan Hang, Janos Demeter, Seung K. Kim, Peter K. Jackson

## Abstract

Multiple G protein coupled receptors (GPCRs) are expressed in pancreatic islet cells but the majority have unknown functions. We observe specific GPCRs localized to primary cilia, a prominent signaling organelle, in pancreatic α- and β-cells. Loss of cilia disrupts β-cell endocrine function, but the molecular drivers are unknown. Using functional expression, we identified multiple GPCRs localized to cilia in mouse and human islet α- and β-cells, including FFAR4, PTGER4, DRD5, ADRB2, KISS1R, and P2RY14. Free fatty acid receptor 4 (FFAR4) and prostaglandin E receptor 4 (PTGER4) agonists stimulate ciliary cAMP signaling and promote glucagon and insulin secretion by α- and β-cell lines, and by mouse and human islets. Transport of GPCRs to primary cilia requires *TULP3*, whose knockdown in primary human and mouse islets depleted ciliary FFAR4 and PTGER4, and impaired regulated glucagon or insulin secretion, without affecting ciliary structure. Our findings provide index evidence that regulated hormone secretion by islet α- and β-cells is regulated by ciliary GPCRs providing new targets for diabetes.

## Introduction

Type 2 diabetes mellitus (T2D) is a pandemic disease affecting over 400 million patients worldwide^1^. Hallmarks of T2D include elevated blood glucose levels, inadequate circulating insulin, and excessive glucagon. Although glucose is a primary mediator of insulin and glucagon release from pancreatic islets, circulating factors including free fatty acids, amino acids, neurotransmitters and hormones like incretins can play critical roles^2^. Accordingly, there is considerable interest in drugs that regulate insulin and glucagon secretion, notably drugs that regulate the G protein coupled receptors (GPCRs) for the incretins Glucagon-like peptide 1 (GLP-1) and glucose-dependent insulinotropic peptide (GIP^3^). Nutrient-sensing GPCRs contribute to many aspects of α- and β-cell function, including regulated insulin and glucagon secretion^2, 4^. Recent studies suggest that primary cilia play critical roles in the β- and alpha cells of the pancreatic islet^5, 6^, but the molecular nature of that function is unclear. Here, we identify specific GPCRs that localize to islet α- and β-cell cilia and regulate glucagon and insulin secretion.

The primary cilium is a membrane and microtubule-based sensory organelle protruding from the apical cell surface and is highly enriched with specialized GPCRs^7^. Transport of GPCRs into the cilium is regulated by two major protein complexes, called the BBSome and the TULP3-IFT-A complex^8^. Defects in primary cilia result in disorders collectively called “ciliopathies”, often present with metabolic syndromes including early onset obesity and eventual diabetes^7^. Cilia are present on mouse and human islet α- and β-cells^9^ and recent evidence has linked diabetic progression in ciliopathy patients to impaired insulin secretion^10–12^. Recent data also showed that dysregulation of cilia associated genes are linked to increased risk of T1D and T2D^13–15^. Moreover, two rodent models have also correlated fewer ciliated α- and β-cells with impaired glucose-regulated insulin secretion^13, 16, 17^.^13, 16, 17^. Moreover, β-cell specific mouse knockouts (KO) of the Ift88 core ciliary gene and Bbs4 component of the BBSome, critical for ciliary trafficking and signaling, ^5, 6, 13^ also show impaired glucose-stimulated insulin secretion. These data argue that cilia are important in the development of diabetes, raising the possibility that ciliary signaling through factors like GPCRs could regulate α- and β-cell function. However, the identity of GPCRs or other signaling regulators that localize to primary cilia and regulate β-cell and α-cell secretion have not been reported.

Here we screened ciliary GPCRs as candidate regulators of insulin or glucagon output by islets. We discovered that GPCRs like FFAR4 and PTGER4, whose natural agonists include omega-3 free fatty acids like DHA and the prostaglandin PGE2, localize to native α- and β-cell cilia, and regulate insulin and glucagon secretion in response to pharmacological agonists through localized ciliary cyclic AMP (cAMP) signaling. We further find that agonists of receptors for omega-3 fatty acids can enhance glucose secretion in response to GLP1R agonists indicating the potential for combination therapies for T2D.

## Results

### Identification of ciliary GPCRs regulating insulin and glucagon secretion

We hypothesized that α- and β-cell ciliary GPCRs transduce signals to regulate islet insulin or glucagon secretion and sought to identify GPCRs that localized to α- and β-cell cilia. From human pancreas transcriptome studies^18^, we identified 96 GPCRs enriched in α- and β-cells compared to pancreatic duct cells (Fig. 1a). To assess ciliary localization, we expressed each candidate GPCR as a C-terminal GFP fusion protein, and assessed subcellular localization in the mouse pancreatic α-cell line α-TC9^19^, and in the mouse β-cell line MIN6^20^, which are both uniformly ciliated (Fig. 1b). We found that FFAR4, PTGER4, ADRB2, KISS1R, and P2RY14 localized to cilia in both MIN6 and α-TC9 cells: (Extended Data Fig. 1a, b). To confirm ciliary localization of candidate GPCRs in native islet cells, we used antibodies that recognize endogenous GPCR protein^21–20^, and found that endogenous free fatty acid receptor 4 (FFAR4) and prostaglandin E receptor 4 (PTGER4) localized to the primary cilium of MIN6 and α-TC9 cells. In contrast, endogenous KISS1R, a receptor for the peptide hormone kisspeptin, was not found localized to cilia in MIN6 and α-TC9 cells. Likewise, endogenous FFAR1, a functional homolog of FFAR4 that also binds omega-3 fatty acids, was not ciliary (Fig. 1c-h; Extended Data Fig. 1c, d). Thus, we successfully identified a critical set of GPCRs that localize to cilia in islet α- and β-cell lines.

**Fig. 1.**
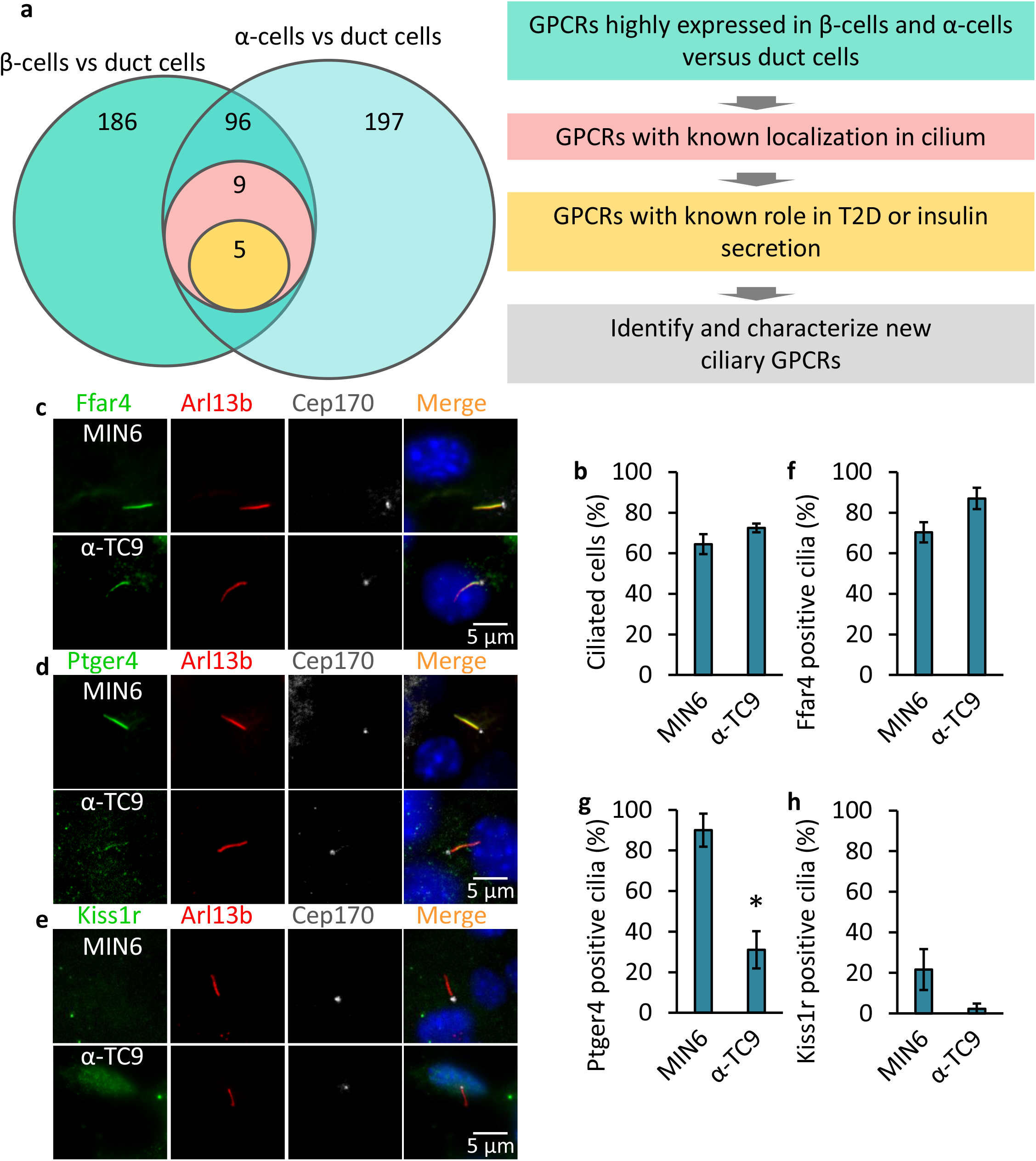
Identification of ciliary GPCRs regulating insulin and glucagon secretion. (**a**) Schematic of the screen to identify ciliary GPCR in human pancreatic α- and β-cells. Candidate GPCRs were selected based on known localization in the cilium and for links to T2D or insulin/glucagon secretion. (**b**) MIN6 and α-TC9 cells are ciliated. MIN6 and α-TC9 cells were grown to confluence. Ciliated cells were examined by confocal fluorescence microscopy using acetylated tubulin and Arl13b antibodies, and quantified. (**c-h**) Endogenous Ffar4 and Ptger4 but not Kiss1r localize to the primary cilium of MIN6 and α-TC9 cells. MIN6 and α-TC9 cells grown to confluence were immunostained with indicated antibodies (**c-e**). Percentages of GPCRs-positive ciliated cells (labelled Ar113b) are shown in **f-h**. Error bars in **b** and **f-h** represent mean ± s.d. (n = 3 independent experiments with 100 cells scored per experiment).

### Ciliary GPCRs regulate insulin and glucagon secretion

To test if FFAR4 or PTGER4 regulate insulin or glucagon secretion, we exposed MIN6 or α-TC9 cells to selective agonists of these GPCRs and measured regulated insulin and glucagon secretion. MIN6 cells displayed glucose-dependent insulin secretion, and α-TC9 cells showed glucose-dependent glucagon secretion, as previously reported^19,20^ (Fig. 2a, b). Next, we examined the effect of agonists for FFAR4 or PTGER4 on insulin or glucagon secretion. Both FFAR4 and PTGER4 agonist treatment augmented insulin secretion in a dose-dependent manner at elevated (16.7 and 25 mM) glucose concentrations compared to controls (Fig. 2c, d). Similarly, the FFAR4 agonist enhanced glucagon secretion at low (1 mM) glucose concentrations in α-TC9 cells (Fig. 2e), whereas a PTGER4 agonist did not (Fig. 2f). We found that agonists of GPCRs not localized to cilia in MIN6 or α-TC9 cells also potentiated glucose-regulated insulin or glucagon secretion. This included exposure to agonists for KISS1-R (Extended Data Fig. 2 a,b), and to TUG424, a selective FFAR1 agonist^22^ (Extended Data Fig. 2 f). In contrast to these results, we observed no detectable effect on MIN6 insulin secretion or α-TC9 glucagon secretion following analogous exposure to agonists of ADRB2 or P2RY14 (Extended Data Fig. 2c, d). Together these data indicate that FFAR4 and PTGER4 are ciliary GPCRs that can potentiate glucose-regulated insulin and glucagon secretion. Glucagon-like peptide 1 receptor (GLP-1R) is a GPCR not known to localize to islet cilia. GLP-1R agonists comprise an important standard treatment for type 2 diabetes^23^. We observed that stimulation of GSIS by the GLP-1R agonist Exendin-4 showed a similar scale of effect to FFAR4 agonists (Extended Data Fig. 2e). To evaluate GLP-1R potentiation of glucose-dependent insulin secretion in the setting FFAR4 activation, we simultaneously stimulated GLP-1R with the agonist Exendin-4, and FFAR4 with TUG891 in MIN6 cells. Compared to exposure to Exendin-4 or TUG891 alone, glucose-dependent insulin secretion by MIN6 cells was substantially increased by simultaneous Exendin-4 and TUG891 (Extended Data Fig. 2e). We observed a similar effect with combined exposure to TUG424, a FFAR1 agonist, and TUG891 (Extended Data Fig. 2f). Thus, FFAR4 agonists could potentially combine with GLP1R and FFAR1 agonists, suggesting these may signal via distinct, but cooperating signaling pathways.

**Fig. 2.**
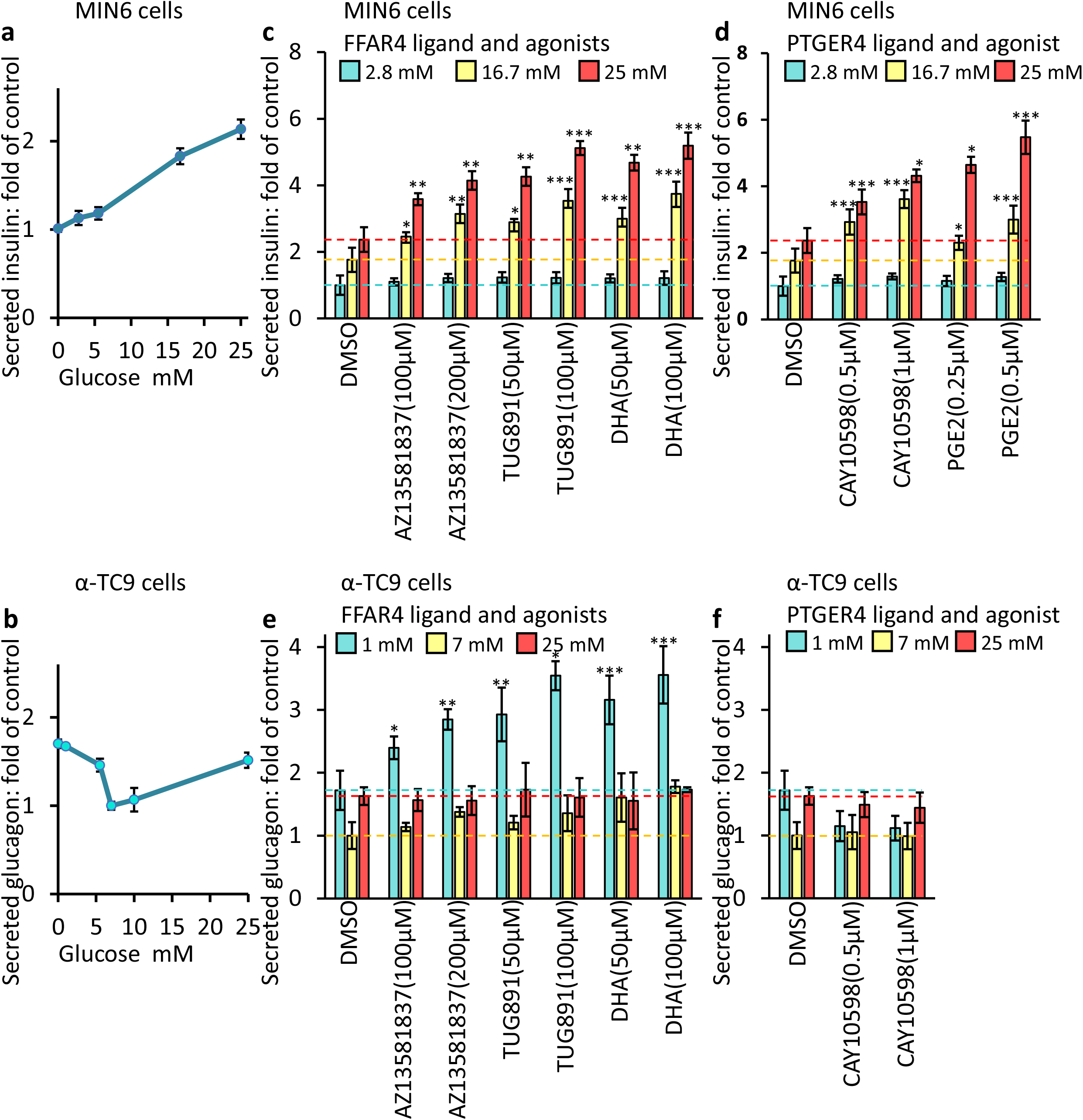
Ciliary GPCRs agonists promote GSIS and GSGS. MIN6 (**a**) and α-TC9 (**b**) cells responded in a dosedependent manner to glucose. GSIS (**c,d**) and GSGS (**e,f**) induced by elevation (from 2.8 mM to 25 mM) or decrease (from 25 mM to 1 mM) of glucose levels and then effects of agonists on insulin or glucagon secretion have been evaluated. Bar graphs are normalized mean ± SD (n = 3 independent experiments); *p < 0.05; **p < 0.01; ***p < 0.001.

### TULP3 is required for trafficking FFAR4 and PTGER4 to islet cell cilia

TULP3 is a crucial regulator of GPCR trafficking and localization to cilia. For example, depletion of TULP3 impairs localization of FFAR4 to the cilium of 3T3L1 preadipocytes and attenuates FFAR4 agonist-regulated adipogenesis^21^. However, prior studies have not assessed the requirement for TULP3 in GPCR trafficking islet cell cilia^8^.To test this possibility, we used CRISPR-Cas9 to generate MIN6 and α-TC9 cells lacking TULP3 (Fig. 3a, b). Consistent with prior work in other cells, depletion of TULP3 from primary cilia in MIN6 and α-TC9 cells did not detectably affect ciliation (Fig. 3c, d). However, TULP3 depletion in these cells clearly reduced localization of the ciliary signaling protein ARL13B, consistent with reports in other cell types (Extended Data Fig. 3a,b)^24^. Moreover, we found that depletion of TULP3 significantly decreased trafficking of FFAR4 and PTGER4 to cilia in MIN6 and α-TC9 cells (Fig. 3e-g). Thus, islet cell ciliary FFAR4 and PTGER4 localization is TULP3-dependent.

**Fig. 3.**
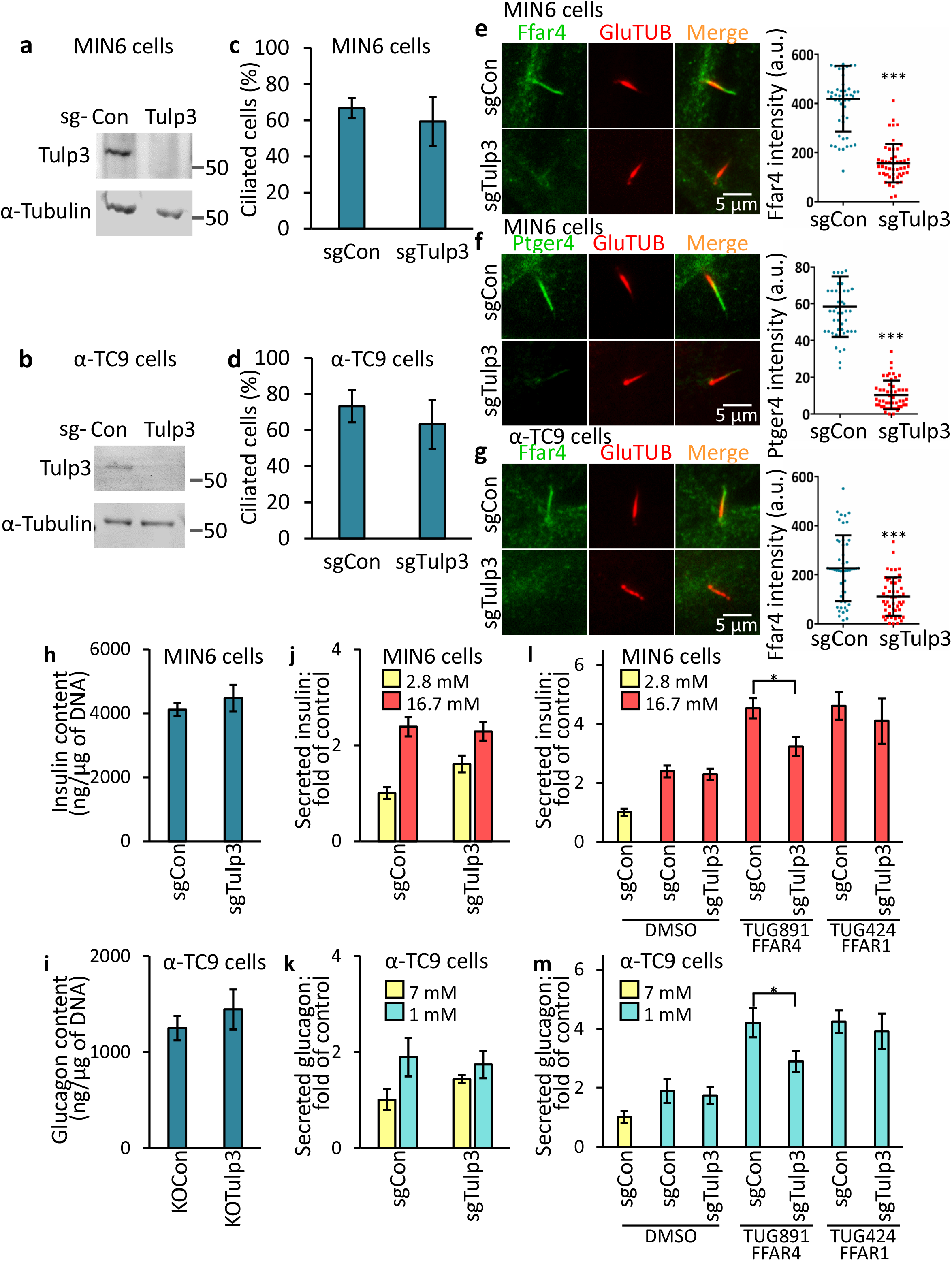
Ffar4-regulated GSIS and GSGS are Tulp3 dependent in pancreatic α- and β-cell lines. (**a-d**) Loss of Tulp3 does not affect cilia formation in MIN6 and α-TC9 cells. Immunoblot showing depletion of TULP3 in MIN6 **(a**) and α-TC9 (**b**) sgTulp3 cell line. Control MIN6 (**c**) and α-TC9 (**d**) cells (sgCon) and Tulp3 knockout cell lines (sgTulp3) grown to confluence. Ciliated cells were examined by confocal fluorescence microscopy using acetylated tubulin antibodie, and quantified. (**e-g**) Loss of Tulp3 prevents ciliary GPCRs trafficking in MIN6 and α-TC9 cells. Control MIN6 (**e,f**) and α-TC9 (**g**) cells and Tulp3 knockout cell lines grown to confluence were immunostained with indicated antibodies. The graph (**e-g**, right) shows the normalized fluorescent intensity of the indicated GPCR proteins that co-localize with the cilium. (**h,i**) Insulin and glucagon content was unchanged in Tulp3 knockout vs control MIN6 (**h**) or α-TC9 (**i**) cells. (**j,k**) GSIS and GSGS was unchanged in Tulp3 knockout vs control MIN6 (**j**) or α-TC9 (**k**) cells. (**l,m**) Ffar4-regulated GSIS (**l**) and GSGS (**m**) are cilia dependent. Error bars in **c**, **d, and h-m** represent mean ± s.d. (n = 3 independent experiments). *p < 0.05; **p < 0.01; ***p < 0.001.

### FFAR4- and PTGER4-regulated insulin or glucagon secretion is cilia dependent

To test whether islet α- and β-cells require ciliary trafficking of FFAR4 and PTGER4 to transduce metabolic cues and potentiate insulin and glucagon secretion, we treated MIN6 or α-TC9 cells lacking TULP3 with the specific FFAR4 and PTGER4 agonists and examined effects on insulin and glucagon secretion. If FFAR4 or PTGER4 regulates insulin or glucagon secretion in a cilia-dependent manner, agonist or ligand treatment should enhance glucose-stimulated insulin or glucagon secretion in control cells but not in cells lacking TULP3. Insulin and glucagon content were not significantly altered in cells lacking TULP3 (Fig. 3h, i). This shows that TULP3 depletion did not affect hormone production or accumulation. TULP3 loss also did not attenuate glucose-dependent insulin or glucagon secretion of MIN6 and α-TC9 cells Fig. 3j, k). However, loss of TULP3 impaired the potentiation by FFAR4 or PTGER4 agonists of glucose-dependent insulin secretion by MIN6 cells (Fig. 3l and Extended Data Fig. 3c) and glucagon secretion by α-TC9 cells (Fig. 3m), arguing that ciliary localization is critical for FFAR4 and PTGER4 signaling in islet cells. By contrast, loss of TULP3 did not reduce potentiation of insulin or glucagon secretion by TUG424, an agonist for FFAR1 which is not cilia-localized (Fig. 3l, m). Likewise, TULP3 knockout did not affect KISS1R-regulated insulin or glucagon secretion (Extended Data Fig. 3c, d). Thus, our work provides evidence that FFAR4 or PTGER4 agonists promote insulin or glucagon secretion via TULP3-dependent mechanisms in primary cilia of MIN6 or α-TC9 cells.

### Localization of FFAR4 and PTGER4 to cilia in primary human and mouse islet cells

Previous studies have shown that human and murine islet cells are ciliated^9, 25^. To confirm and extend these observations, we first quantified ciliation of islet α-, β-, and δ-cells from mice or human cadaveric donors and found that both mouse and human endocrine cells are efficiently ciliated, about 62, 70, and 75% in mouse and about 32, 41, and 33% in human α-, β-, and δ-cells (Fig. 4a, b). We noted no significant difference in the percentage of ciliation of these islet cell subsets in mice versus humans. Supporting our findings with α- and β-cell lines, we found that FFAR4 localized to the primary cilium of pancreatic α- and β-cells in mouse and human islets, and that this ciliary localization persisted even after dissociation and flow cytometry-based purification of α- and β-cells from isolated islets (Fig. 4c-f and Extended Data Fig. 4a, b). PTGER4 also localized to the primary cilium in β-cells, but compared to FFAR4, the degree of PTGER4 localization to α-cell cilia was lower (Fig. 4g-j and Extended Data Fig. 4c, d). We next examined the effects of FFAR4, PTGER4, and KISS1R agonists on insulin or glucagon secretion from mouse or human islets. Addition of agonists promoted insulin or glucagon secretion in mouse or human islets (Extended Data Fig. 4g-j). Consistent with our cell line data, PTGER4 agonist did not affect glucagon secretion (Extended Data Fig. 4h, j). Taken together, our data shows that FFAR4 and PTGER4 localize to the cilium in mouse and human endocrine cells and regulate insulin and glucagon secretion.

**Fig. 4.**
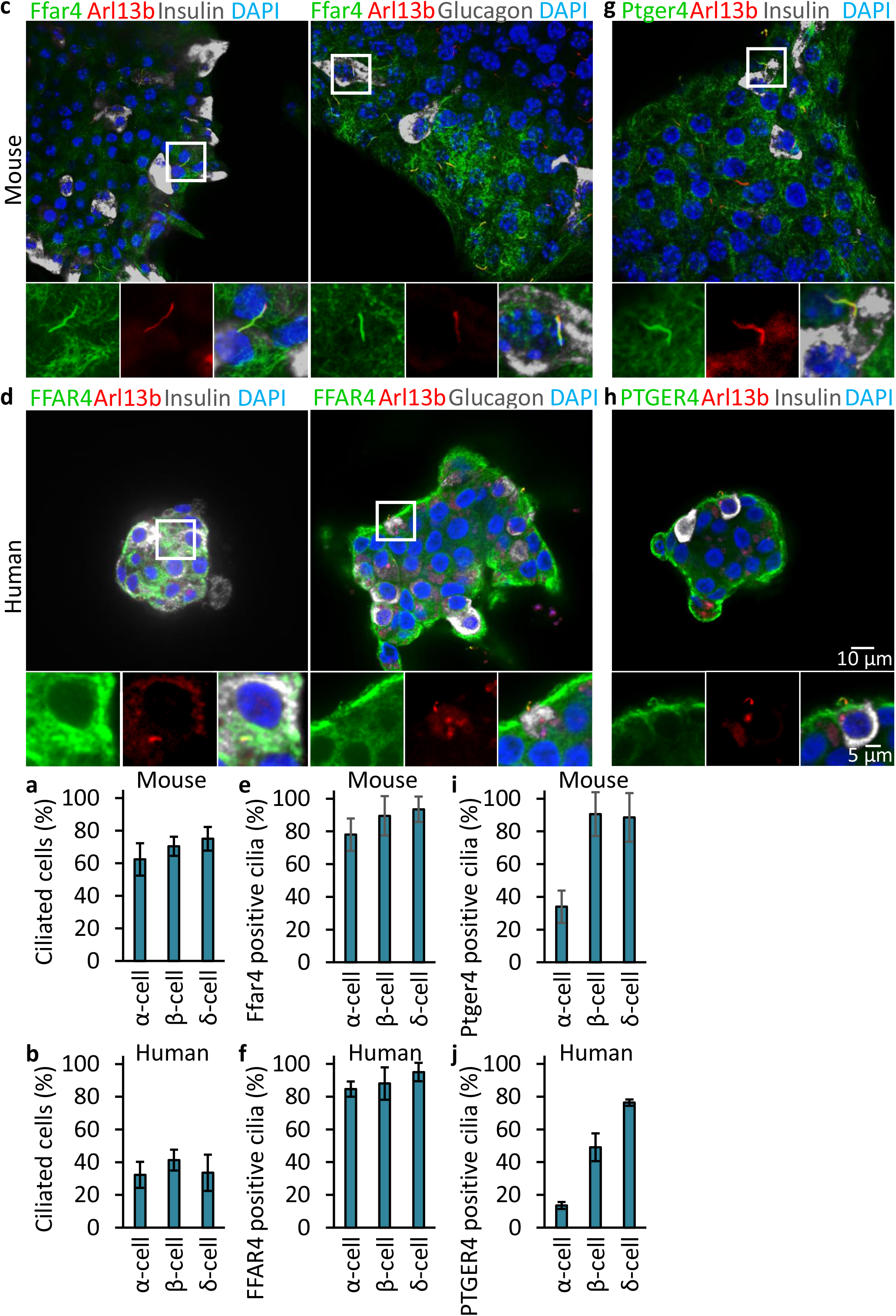
FFAR4- and PTGER4 are a ciliary GPCR displayed by mouse and human pancreatic α- and β-cells. (**a,b**) α-, β-, and δ-cells are ciliated in mouse (**a**) and human (**b**) islet. Ciliated cells were examined by confocal fluorescence microscopy using acetylated tubulin, Arl13b, insulin, glucagon, and somatostatin antibodies, and quantified. (**c-j**) Endogenous Ffar4 and Ptger4 localize to the primary cilium of α- and β-cells in mouse (**c, g, e, i**) and human (**d, h, f, j**) pancreatic islet. (**e, f, i, j**) Percentages of GPCRs-positive mouse (**e, i**) and human (**f, j**) ciliated α-, β-, and δ-cells (labelled Ar113b) are shown. Error bars in **a–j** represent mean ± s.d. (n = 3 independent experiments with 100 cells scored per experiment).

### FFAR4- and PTGER4-regulated islet hormone secretion is cilia dependent

To test if FFAR4- and PTGER4-regulated hormone secretion in primary pancreatic islets requires *TULP3*, we infected dispersed primary human islet cells with lentivirus expressing shRNA to knockdown *TULP3*, re-aggregated these cells into pseudoislets^26^, and measured glucose-dependent insulin or glucagon secretion. Lentiviral GFP co-expression permitted sorting and isolation of virus-infected cells. >50% knockdown of *TULP3* mRNA was demonstrated by qRT-PCR (Extended Data Fig. 5a, b). Total insulin or glucagon content was not significantly altered after *TULP3* knockdown (Extended Data Fig. 5c-f). Similar to our islet cell line studies, depletion of *TULP3* mRNA did not affect insulin or glucagon secretion in response to glucose changes alone (Fig. 5a, c, e, g), but did significantly attenuate FFAR4 and PTGER4 agonist-regulated insulin or glucagon secretion (Fig. 5b, d, f, h). Similar to our cell line studies, exposure to selective FFAR1 and KISS1R agonists (whose receptors do not localize to islet cilia) also stimulated insulin or glucagon secretion, and this effect was not altered by *TULP3* islet knockdown (Fig. 5b, d, f, h and Extended Data Fig. 5g-j). Thus, our findings establish that FFAR4- and PTGER4-regulated insulin or glucagon secretion by primary islet cells are TULP3-dependent and function efficiently in mouse and human islets.

**Fig. 5.**
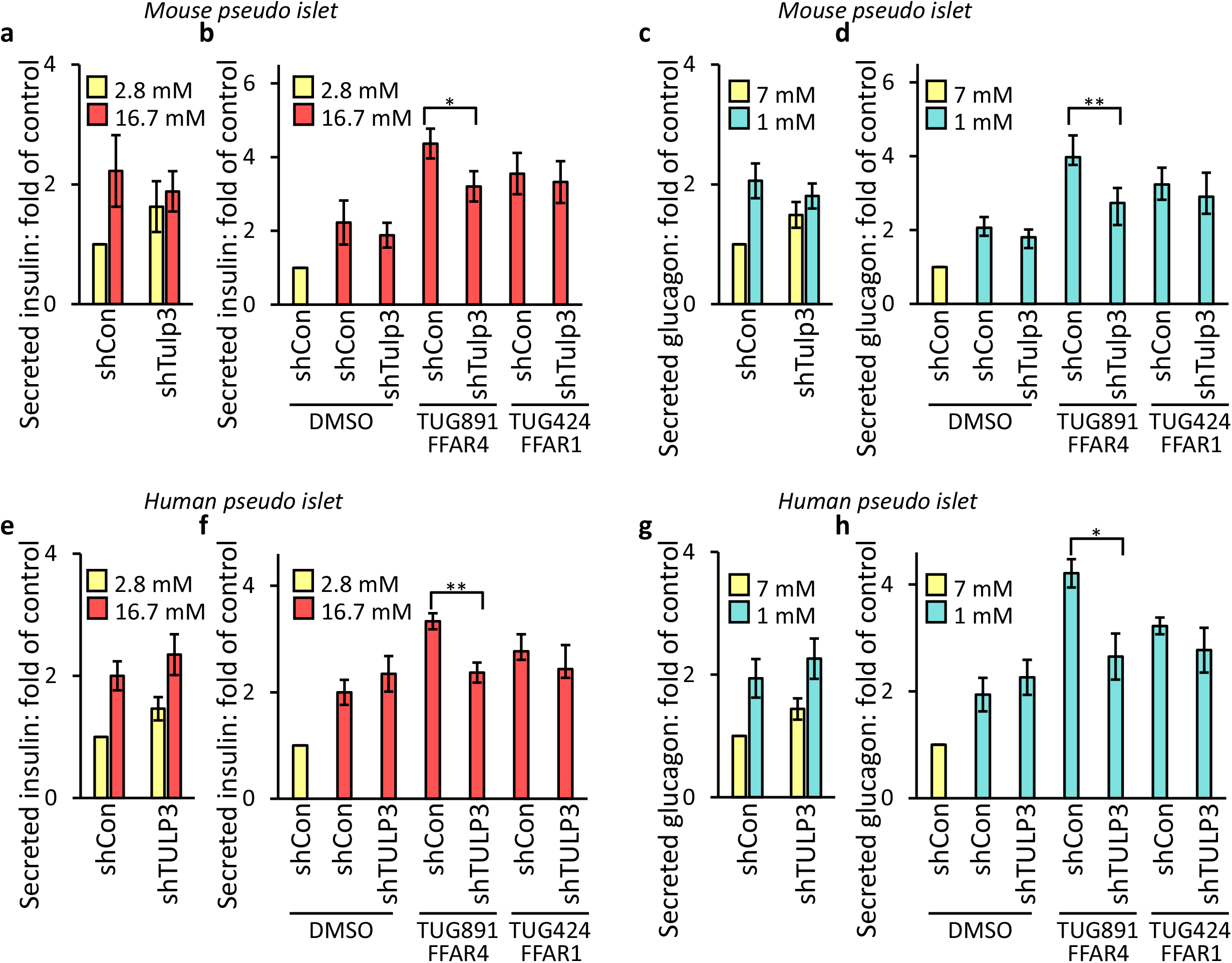
FFAR4-regulated GSIS and GSGS are cilia dependent in pancreatic islet. GSIS (**a, e**) and GSGS (**c, g**) was unchanged in Tulp3 knockdown vs control mouse (**a,c**) or human (**e, g**) pseudoislets. (**b, d, f, h**) Ffar4-regulated GSIS (**b, f**) and GSGS (**d, h**) are cilia dependent in mouse (**b, d**) and human (**f, h**) pseudoislets. Error bars in **a-h** represent mean ± s.d. (n = 4 independent experiments). *p < 0.05; **p < 0.01; ***p < 0.001.

### FFAR4 signaling raises ciliary cAMP to promote insulin or glucagon secretion

To identify molecular signaling mechanisms underlying FFAR4 or PTGER4 potentiation of insulin or glucagon secretion, we investigated ciliary cAMP signaling, which was shown in our recent studies to mediate omega-3 fatty acid activation of ciliary FFAR4, and to trigger adipogenesis^21^. Adenylyl cyclases (ACs) catalyze the conversion of adenine triphosphate (ATP) into cAMP in response to a wide range of extracellular signals. In mammals, there are nine membrane-associated ACs. The type 3 adenylyl cyclase (AC3), an established cilia marker, is highly and predominantly expressed in primary cilia in different tissues, including pancreas, adipose tissue, kidney and brain^27^. Moreover, recent human studies show that mutation of the *ADCY3* gene encoding AC3 is associated with T1D and T2D^14, 15^. We observed that AC3 localizes to MIN6 and α-TC9 cilia (Extended Data Fig. 6a). To test the hypothesis that FFAR4 activation promotes insulin or glucagon secretion by modulating AC3-regulated ciliary cAMP, we transduced MIN6 and α-TC9 cells with a ciliary-targeted cAMP sensor optimized for live cell imaging (cilia cADDIS^28^). Within 120 seconds of MIN6 or α-TC9 exposure to the FFAR4 agonist TUG891, we observed increased ciliary cAMP levels (Fig. 6a, b and Extended Data Fig. 6b). As a control, a selective FFAR1 agonist did not potentiate cAMP levels. cAMP may activate at least two downstream pathways regulated by protein kinase A (PKA) or by the cAMP-regulated exchange factor, EPAC. Inhibition of EPAC prevented FFAR4 agonist-stimulated insulin and glucagon secretion (Fig. 6c, d). Inhibition of PKA also inhibited FFAR4 agonist-stimulated insulin secretion but not glucagon secretion (Fig. 6e, f). Thus, FFAR4 signaling likely potentiates insulin secretion by activating localized ciliary cAMP signaling via both EPAC and PKA activity. Interestingly, prior studies show that FFAR4 agonist regulated adipogenesis through activation of localized ciliary cAMP and EPAC, but not detectably via PKA^21^. In studies examining potentiation of secretion, we found that inhibition of EPAC, but not PKA, prevented PTGER4 agonist-enhanced insulin secretion (Fig. 6c, e). Together these studies identify cAMP-dependent mechanisms of ciliary GPCR signaling. This suggests that signal transduction through specific GPCRs is executed through overlapping but distinct pathways.

**Fig. 6.**
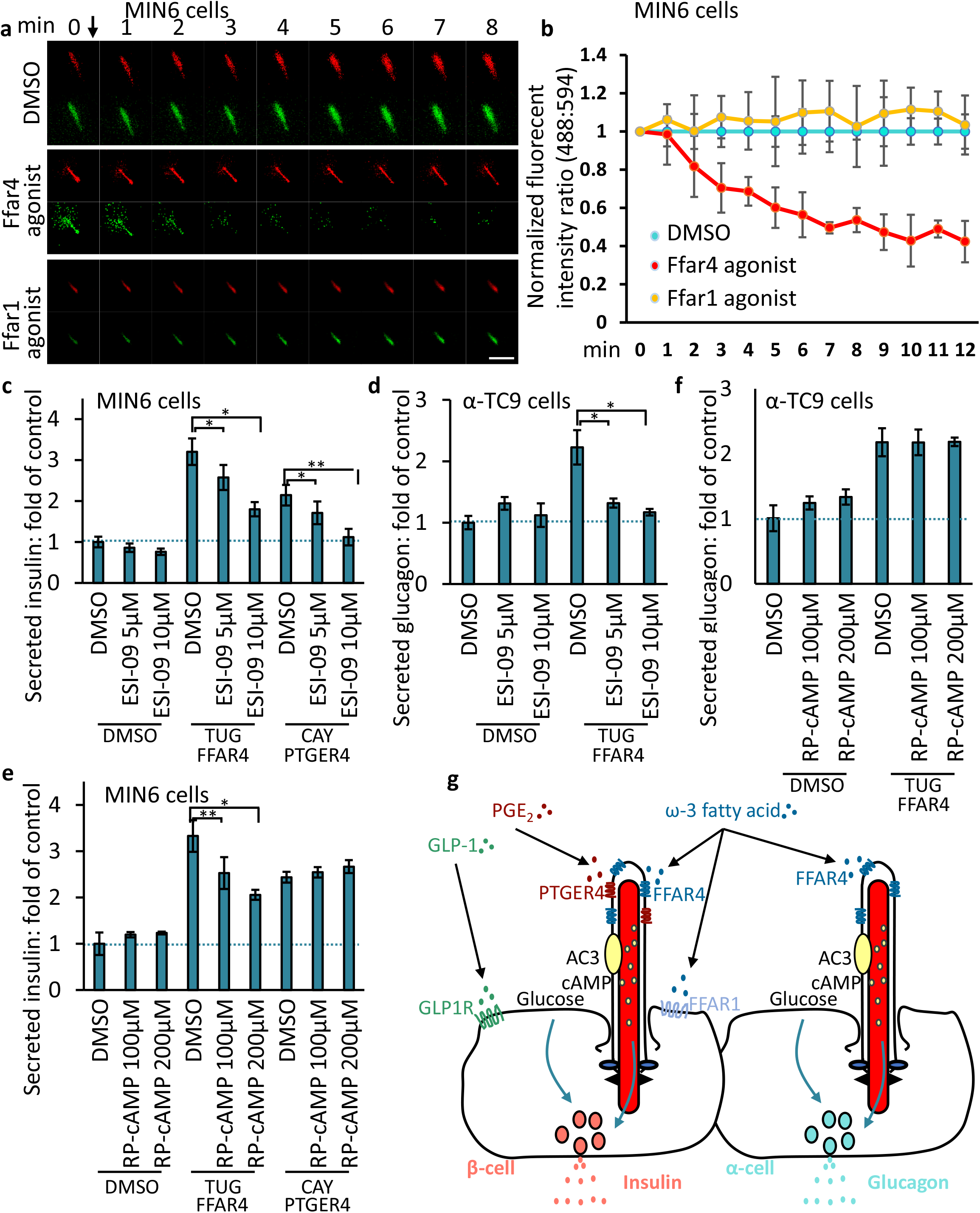
FFAR4 Regulates GSIS and GSGS via cAMP. (**a**) Representative images showing the cADDIS cAMP sensor (green) and cilia (red) offset in MIN6 cells. Scale bar, 2 mm. (**b**) Background subtracted ratio of fluorescence intensities are normalized to DMSO control and 0 s time point. n = 3 for FFAR4 agonist and DMSO control ± SD, where n is the average of all cilia measured per well. (**c-f**) Inhibition of EPAC attenuates Ffar4-regulated GSIS and GSGS in a dosedependent manner. MIN6 (**c, e**) and α-TC9 (**d, f**) cells were stimulated with GPCR agonists in the presence of an inhibitor of EPAC (ESI-09) or PKA (Rp-cAMPS) for the first 1 h. Error bars in **c–f** represent mean ± s.d. (n = 3 independent experiments). *p < 0.05; **p < 0.01; ***p < 0.001. (**g**) Model for ciliary GPCR-regulated insulin or glucagon secretion. FFAR4 localized to the cilia in α- and β-cell is activated by ω-3 fatty acids to promote glucose-stimulated insulin or glucagon secretion. PTGER4 localized to cilia in β-cells is activated by PGE_2_ to regulate glucose-stimulated insulin secretion. GLP1R and FFAR1 localized to the plasma membrane are activated by GLP-1 or ω-3 fatty acid to regulate insulin secretion, which can cooperate with FFAR4 stimulated insulin secretion.

## Discussion

Pancreatic islet cells are vital regulators of metabolism that integrate diverse, dynamic signals, to optimize their output of insulin and glucagon. While glucose is a primary regulator of insulin and glucagon secretion by islets, other important signals reflecting metabolic flux or feeding state are recognized as crucial controllers of islet hormone output. Thus, multiplexed signaling from the autonomic and central nervous system, peripheral organs like fat, liver, inflammatory cells and peripheral endocrine organs, and dietary sources tunes islet hormonal output to match dynamic physiological demands in healthy and diseased states ^29, 30^. Intensive investigations have focused on identifying the cellular and molecular signaling elements in islets that integrate and orchestrate islet hormone output^31^. Work here provides index evidence that islet cell primary cilia are organizers of signaling by specific G-protein coupled receptors (GPCRs) that respond to native ligands and synthetic agonists to regulate islet insulin and glucagon secretion. Findings here also delineate specific intracellular responses to ciliary GPCR activation and suggest how ciliary signaling logic could integrate cues to optimize hormonal output and metabolic control.

Our work reveals multiple ciliary GPCRs as conserved regulators of insulin and glucagon output by mouse and human islets. Depletion of TULP3, a regulator of GPCR trafficking to cilia in pancreatic α- and β-cells provided crucial, index evidence that ciliary localization of GPCRs like FFAR4 and PTGER4 is required for potentiation of insulin or glucagon secretion by specific agonists *in vitro* and *ex vivo*. Agonist activation of ciliary FFAR4 and PTGER4 resulted in a rapid increase in cAMP levels, activating EPAC, a guanine-nucleotide exchange factor, and protein kinase A. Moreover, demonstration of signaling effects in α and β cell lines argues that TULP3-dependent ciliary GPCR effects are cell-autonomous (Fig. 6g). Our studies provide critical functional evidence for how ciliary GPCR signaling works in islet cells and identifies candidate small molecules that regulate insulin or glucagon secretion.

This study provides evidence that ciliary FFAR4 and PTGER4 regulated insulin secretion by β-cells. In addition, we show that ciliary FFAR4 regulates glucagon secretion by α-cells^32^. Specifically, *in vivo* and *ex vivo* loss of ciliary FFAR4 in α- and β-cells impairs FFAR4 agonist-regulated insulin and glucagon secretion. We also found that this process is dependent on ciliary cAMP (Fig. 6g). While cAMP is generally considered as an amplifier of insulin secretion triggered by Ca^2+^ elevation in the β-cells^33^ it was not previously reported that subcellular organelle signaling, like in islet cell cilia, might underlie this effect. cAMP is synthesized by adenylyl cyclases (ACs). In mammals, there are nine membrane-associated ACs. Here, we found that AC3 localizes to the cilia in MIN6 and α-TC9 cells. This indicates that ciliary AC3 may be activated by FFAR4 and then increase the cAMP level in the cilium. However, we recognize other ACs like AC5/6 may also be involved; recent data showed that mutation of ADCY3, ADCY5, and ADCY6 are each strongly associated with T1/T2D^14, 15, 34, 35^. We hypothesize that specific cAMP effectors may work locally at cilia or centrioles to control signaling or transport processes. In many organisms including unicellular protists, primary cilia or flagella are localized next to zone of organized endocytosis and exocytosis. The specific configuration of cilia in the context of islet cells is therefore of interest, and future studies could address how localized cAMP signals are propagated to the secretory structures in islet cells. In addition to signaling mechanisms revealed here, prior reports^36^ suggest that FFAR4 stimulation may lead to activation of phospholipase C (PLC) and subsequent elevation of intracellular Ca^2^; we have found that exposure of MIN6 and α-TC9 cells to PLC inhibitor U73122 also attenuates FFAR4-stimulated insulin and glucagon secretion (data not shown). Thus, additional studies will also be useful for understanding how the same ciliary GPCR can simultaneously and locally modulate multiple signaling pathways, and whether this multiplex signaling is achieved by dynamic control of localized intra-or juxta-ciliary signaling.

PGE_2_ production is activated in response to various forms of pancreatic damage, and is expected to work via four independent receptors PTGER1-4, often called EP1-4. Of these four, only PTGER4/EP4 is ciliary^37^ (PKJ, unpublished) and is the only form linked to Gα_s_ and cAMP production. Prior studies have shown a protective role for EP4 on β cell survival and proliferation^38^, but EP3 may have opposing effects. The presence of inflammatory receptors like PGTER4 may couple localized pancreatic or more global responses to islet output of insulin. Systemic delivery of high levels of PGE2 appear to lower insulin secretion^39, 40^, but the selective effects on β cell proliferation or survival versus direct effects on insulin secretion are not fully separated. The broader context of activating PGE_2_ levels in pancreatic islets may cause a variety of effects. Here, by using selective EP4 agonists, we show that direct effects on cAMP, via its ciliary signaling channel, have a positive effect on insulin release, supporting the potential value of using selective drugs for EP4. This highlights the potential importance of selective signaling through multiple islet ciliary GPCRs as a mechanism to organize responses by a single cell. By organizing islet cell sensors, notably in the form of G-protein coupled receptors, the multicellular islet cluster integrates and tailors the potent systemic response of islet hormones to systemic demands.

Our work also revealed evidence of additive or synergistic regulation of β cell insulin secretion by FFAR4 and FFAR1, or by FFAR4 and GLP1-R (Extended Data Fig. 2e, f). This is supported by recent studies suggesting that FFAR1 and FFAR4 signaling can cooperate to stimulate insulin secretion^41^. Neither FFAR1 (Fig. 1) nor GLP1-R (CT.W. and P.J., unpublished results) are observed to localize to cilia. Consistent with this, FFAR1-regulated insulin and glucagon secretion is TULP3 independent. Thus, outcomes here suggest that islet hormone secretion may be regulated by simultaneous activation of GPCR-dependent signaling pathways in *distinct* subcellular compartments (Fig. 6g). FFAR1 and FFAR4 bind to medium-to long-chain fatty acids, including ω-3 fatty acids like α-linolenic acid, eicosapentaenoic acid (EPA), and docosahexaenoic acid (DHA), leading to enhanced insulin and glucagon secretion^22^, although it is not entirely clear that FFAR1 and FFAR4 signaling results from precisely the same natural ligands, Unlike FFAR4 agonists, FFAR1 agonists do not raise ciliary cAMP, but instead stimulate calcium influx^41^. We also observe evidence of synergy between FFAR4 agonists and GLP1R agonists (Extended Data Fig. 2e, f). GLP1R signaling can induce intracellular Ca^2+^ transients in addition to cAMP signaling^42^. Thus, if ciliary FFAR4-cAMP signaling were to activate docked, active zone vesicles, possibly through EPAC2-dependent signaling, this could augment or synergize with GLP1-R or FFAR1-calcium dependent insulin secretion.

Based on the signaling we observe here in isolated islets, we can propose that the apical position of primary cilia in pancreatic islets provide a critical architecture that integrates islet cell cross-talk. This could represent both homotypic (β cell to β cell) and heterotypic (β cell to α cell) paracrine signals from nearby islet cells, but also global endocrine and metabolite signals. We can imagine a range of signaling inputs, some commonly used and some more specialized, that would control insulin and glucagon secretion, and more broadly in other islet subtypes. There is the intriguing possibility that ciliary signaling in multiple islet cell types allows an integration of dietary and neuroendocrine signals, ensuring metabolic homeostasis and rapid responses to specialized conditions.

The integrated, multisystem approach used here to identify ciliary GPCRs and mechanisms that regulate human insulin and glucagon secretion can be readily expanded. Further studies could take advantage of the tools and approaches generated here, including development of *Tulp3*-deficient a and b cell lines, pseudoislet-based genetic methods for generating primary human islets lacking TULP3, and measures of agonist-dependent hormone secretion. In addition to GPCRs, other classes of receptors and their signal transduction elements could be revealed by these future studies. In addition, the increasing availability of cadaveric human islets from donors with specific pathological or physiological states could broaden the impact of our findings. For example, based on prior studies, dynamic physiological changes like sexual maturation or pregnancy^43^, or inflammatory states like diabetes^44^ could regulating or dysregulate islet ciliary signaling. If so, this could advance the concept that age or disease-dependent degeneration of ciliary signaling might underlie diabetes and other human pancreatic diseases.

## Acknowledgments

The authors acknowledge members of the Kim laboratory, especially Jonathan Lam and Dr. Sangbin Park for helpful discussions and assistance with islet experiments. MIN6 cells were gifts from Professor Jun-ichi Miyazaki (Department of Stem Cell Regulation Research, Graduate School of Medicine, Osaka University, Osaka, Japan). We thank the Alberta Diabetes Institute Islet (ADI) Research Core, IIDP, NDRI and IIAM for islet and/or pancreas procurement, and especially the organ donors and their families.

## Funding

P.K.J. was supported by NIH grants R01GM11427604, R01HD085901, R01GM12156503 and the Stanford Department of Research, Baxter Laboratory and a Stanford Diabetes Research Center (SDRC) Pilot and Feasibility Research Grant (to P.K.J and S.K.). K.I.H. is a Layton Family Fellow of the Damon Runyon Cancer Research Foundation (DRG-2210-14). R.B. was supported by a postdoctoral fellowship from JDRF (3-PDF-2018-584-A-N). Work in the Kim group was supported by NIH awards (R01 DK107507; R01 DK108817; U01 DK123743; R01DK126482 to S.K.K.), gift funding from Michelle and Steve Kirsch, the Reid family, the Schaffer family fund, the Snyder Foundation, two anonymous donors, and the JDRF Center of Excellence (to S.K.K. and M. Hebrok). Work here was also supported by NIH grant P30 DK116074 (S.K.K.), and by the Stanford Islet Research Core, and Diabetes Genomics and Analysis Core of the Stanford Diabetes Research Center.

## Methods

### Human islet procurement

De-identified human pancreatic islets were obtained from organ donors without a history of glucose intolerance with less than 15 h of cold ischemia time. Islets were procured through the Integrated Islet Distribution Program, Alberta Diabetes Institute IsletCore, and the International Institute for the Advancement of Medicine.

### Mouse islet isolation

Islets were isolated from male C57BL/6 mice at 2 to 4 month of age using Collagenase P (Roche Diagnostics, catalog number: 11213865001) into the pancreatic duct, surgically removing the infused pancreas and placing it into 50-ml conical tubes containing 4 ml of HBSS/Ca/HEPES solution (1 L of Hanks balanced salt solution (HBSS), 2 mM CaCl2, and 20 mM HEPES). Mouse pancreata were incubated for 12 min in a 37 °C water bath. The digested pancreata were than washed three times in ice-cold HBSS/Ca/HEPES solution. The pancreas tissue was disrupted by vigorously hand shaking the tubes for 1 min at a rate of 3 shakes per second. The islets were isolated from acinar tissue on a Histopaque-1077 gradient (Sigma-Aldrich, catalog number: H8889). After three additional washes by RPMI 1640 (Gibco), islets were handpicked under a dissecting microscope and cultured in RPMI 1640, 2.25 g/dl glucose, 1% penicillin/streptomycin (v/v, Gibco) and 10% fetal bovine serum (HyClone).

### Human and mouse pseudo islet generation

Human or mouse islets were dissociated into a single cell suspension by enzymatic digestion (Accumax, Invitrogen). For each experimental condition, ~1 × 10^6^ cells were transduced with lentivirus corresponding to 1 × 10^9^ viral units in 1 ml as determined by the Lenti-X qRT-PCR titration kit (Clonetech). Lentiviral transduced islets cells were cultured in 96-well ultra-low attachment plates (Corning) and cultured for 3 days at 37 °C in 5% CO^2^. After 3 days, pseudo islets were transferred to a 6 well ultra-low attachment plates and cultured 2 days prior to further molecular or physiological analysis. The islets were cultured in culture media: RPMI 1640 (Gibco), 2.25 g/dl glucose, 1% penicillin/streptomycin (v/v, Gibco) and 10% fetal bovine serum (HyClone).

### Cell line models

Pancreatic MIN6 β cells (passages 5–15) were a gift from Professor Jun-ichi Miyazaki (Department of Stem Cell Regulation Research, Graduate School of Medicine, Osaka University, Osaka, Japan). MIN6 and α-TC9 cells were cultured in DMEM medium containing 10% Fetal Bovine Serum, HEPES (1M), 2-Mercaptoethanol (50mM), 1% Pen/Strep, and 1% GlutaMAX.

### Lentivirus production

Lentiviruses were produced by transient transfection of HEK293T cells with lentiviral vectors carrying the gene of interest and pMD2.G (12259; Addgene) and psPAX2 (12260; Addgene) packaging constructs. DMEM Media was re-placed after overnight and virus was harvested 24, 48, and 72 h post-transfection. Virus was filtered with a 0.45mm OVDF filter (Millipore). Supernatants were collected and purified using PEG-it (System Biosciences). Concentrated lentivirus was stored at −80 °C for transduction of primary human cells.

### Cell line generation

Virus carrying the gene of interest was used to infect cell lines with 10mg/mL polybrene (Millipore). Media was replaced after 24 h and cells were sorted for GFP positivity after 48-72 h post-infection. To generate Crispr/Cas9 knockout cells, MIN6 and α-TC9 Cas9-BFP cells were infected with lentivirus containing the sgRNA of interest. Knockout efficiency was determined 10 days post-infection by western blotting. MIN6 and α-TC9 cells expressing Cas9-BFP were generated by infection of virus harvested from 293T cells transfected with p293 Cas9-BFP, pMD2.G and psPAX2. MIN6 and α-TC9 Cas9-BFP cells were sorted for BFP positivity.

### Immunofluorescence staining

Cells were grown on 12mm round coverslips and fixed with 4% paraformaldehyde (433689M, AlfaAesar) in PBS at room temperature for 10min. Samples were blocked with 5% normal donkey serum (017-000-121, Jackson ImmunoResearch) in IF buffer (for FFAR4 staining: 3% BSA and 0.4% saponin in PBS; for all else: 3% BSA and 0.1% NP-40 in PBS) at room temperature for 30min. Samples were incubated with primary antibody in IF buffer at room temperature for 1 h, followed by 5 washes with IF buffer. Samples were incubated with fluorescent-labeled secondary antibody at room temperature for 30min, followed by a 5 min incubation with 4’,6-dia-midino-2-phenylindole (DAPI) in PBS at room temperature for 5min and 5 washes with IF buffer. Coverslips were mounted with Fluoromount-G (0100-01, SouthernBiotech) onto glass slides followed by image acquisition. Antibodies were used as follows: FFAR4 (residues PILYNMSLFRNEWRK, 1:600)^21^, FFAR4 (Santa Cruz, sc-390752, 1:100), Acetylated tubulin (Sigma, T7451, 1:2000), PTGER4 (Santa Cruz, sc-55596, 1:100), KISS1R (A kindly gift from Professor Kirk Mykytyn, The Ohio State University, 1;500), ARL13B (UC Davis/NIH NeuroMab Facility, 73-287, 1:1000), FGFR1OP (Novus,H00011116-M01, 1:1000), GFP (Invitrogen, A10262, 1:2000).

### Epi-fluorescence and confocal imaging

Images were acquired on an Everest deconvolution workstation (Intelligent Imaging Innovations) equipped with a Zeiss AxioImagerZ1 microscope and a CoolSnapHQ cooled CCD camera (Roper Scientific) and a 40x NA1.3 Plan-Apochromat objective lens (420762-9800, Zeiss) was used. Confocal images were acquired on a Marianas spinning disk confocal (SDC) microscopy (Intelligent Imaging Innovations). For Figures 2 and S2, images were acquired using a Leica DMi8 microscope equipped with a DFC7000T color camera (bright field images) as well as the SPE confocal system (immunofluorescence).

### Sample preparation and immunoblot

Cells were lysed in 1x LDS buffer containing DTT and incubated at 95°C for 20 min. Cells were lysed in RIPA buffer with protease inhibitors (50 mM Tris-HCl, pH 8.0, 150 mM NaCl, 1% NP-40, 20 mM β-glycerophosphate, 20 mM NaF, 1 mM Na3VO4, and protease inhibitors including 1 μg/μl leupeptin, 1 μg/μl pepstatin, and 1 μg/μl aprotinin) for 30 min at 4°C. The cell lysates were centrifuged at 16,000 g at 4°C for 15 min. Proteins were separated using NuPage 4%–12% Bis-Tris gel (Thermo Fisher Scientific, WG1402BOX) in NuPage MOPS SDS running buffer (50 mM MOPS, 50 mM TrisBase, 0.1% SDS, 1 mM EDTA, pH 7.7), followed by transfer onto PVDF membranes (Millipore, IPFL85R) in transfer buffer (25 mM Tris, 192 mM glycine, pH 8.3) containing 10% methanol. Membranes were blocked in non-fat dry milk in PBS for 30 min at room temperature, followed by incubation with primary antibody in blocking buffer for overnight at 4°C. The membrane was washed 4 times for 10min in TBST buffer (20 mM Tris, 150 mM NaCl, 0.1% Tween 20, pH7.5) at room temperature, incubated with secondary IRDye antibodies (LI-COR) in blocking buffer for 1 h at room temperature, and then washed 4 times for 10min in TBST buffer. Membranes were scanned on an Odyssey CLx Imaging System (LI-COR), with protein detection at 680 and 800 nm. Antibodies were used as follows: TULP3 (Yenzym, 1:2000)^21^, Tubulin (Sigma, 9026, 1:5000).

### Quantitative Real time PCR

RNA was extracted using the RNeasy Lipid Tissue Kit (QIAGEN) and cDNA was synthesized using M-MLV Reverse Transcriptase (Invitrogen, 28025-013). Quantitative real time PCR was performed using TaqMan Probes (Invitrogen) and the TaqMan Gene Expression Master Mix (Applied Biosystems, 4369016) in 96-well Micro Amp Optical reaction plates (Applied Biosystems, N8010560). Expression levels were normalized to the average expression of the housekeeping gene.

### Live Cell Ciliary cAMP assay

MIN6 and α-TC9 cells were seeded at 10^4^ cells/well in a 96-well cell imaging plate (Eppendorf, 0030741013) and transduced the following day with the ratiometric cilia-targeted cADDis BacMam (Molecular Montana, D0211G) according to manufacturer’s recommendation. Briefly, cells were infected with 25ul of BacMam sensor stock in a total of 150ul of media containing 2mM Sodium Butyrate (Molecular Montana) for overnight in the 37 °C incubator. Prior to imaging, cells were incubated in PBS for 30min at room temperature. Images were acquired on a Marianas spinning disk confocal (SDC) microscopy (Intelligent Imaging Innovations) (40x, epi-fluorescence) every 1min for 15min with agonist added after 30sec. Red fluorescence was used to determine a mask and background subtracted green and red fluorescent intensity over time was determined using Slidebook (Intelligent Imaging Innovations).

### In vitro insulin and glucagon secretion assays

MIN6 and α-TC9 (1×10^5^) seeded in 96-well plates and batches of 25 pseudo islets were used for in vitro secretion assays. MIN6, α-TC9 cells, and pseudo islets were incubated at a glucose concentration of 2.8 mM for 60 min as an initial equilibration period. Subsequently, MIN6, α-TC9 cells, and pseudo islets were incubated at 2.8 mM, 16.7 mM and 16.7 mM + agonists glucose concentrations for 60 min each. Pseudo islets were then lysed in an acid-ethanol solution (1.5% HCL in 75% ethanol) to extract the total cellular insulin or glucagon content. Secreted human insulin or glucagon in the supernatants and pseudo islet lysates were quantified using either a human insulin ELISA kit or glucagon ELISA kit (both from Mercodia). Secreted insulin levels were divided by total insulin content and presented as a percentage of total insulin content; a similar method of data analysis was employed for glucagon secretion assays. All secretion assays were carried out in RPMI 1640 (Gibco) supplemented with 2% fetal bovine serum (HyClone) and the above-mentioned glucose concentrations.

### Reagents and treatment

The concentration of the following reagents was indicated in the figure legend. ESI-09, RP-cAMP, Compound 19a (CAY10598), Salbutamol, UDP-α-D-Glucose and Exendin-4 were from Cayman Chemical. AZ13581837 purchased from AOBIOUS. TUG891 and TG424 were from Tocris. Small molecules were dissolved in DMSO (276855, Sigma-Aldrich). kisspeptin-10 purchased from Santa Cruz. Small molecules or peptides were dissolved in DMSO (276855, Sigma-Aldrich). ESI-09 or RP-Camp 30 min prior to agonist or vehicle addition. The following reagents were used at the indicated concentrations in Fig. 3, Extended Data Fig. 3, Fig. 5, Extended Data Fig. 5, and Fig. 6: 100μM TUG891, 1μM CAY10598, 2μM KP-10, 100nM TUG424.

### Quantification and statistical analysis

Statistical analyses were performed in Microsoft Excel and GraphPad Prism. Most data are represented as mean ± standard derivation (s.d.) as specified in the figure legends. Sample size and number of repeated experiments are described in the legends. p value was determined using the two-tailed unpaired Student’s t-test. The precise P values are shown in the figures. P < 0.05 was considered statistically significant. All experiments were repeated three or four times (see figure legends) with similar results.

---

**Extended Data Fig. 1.**
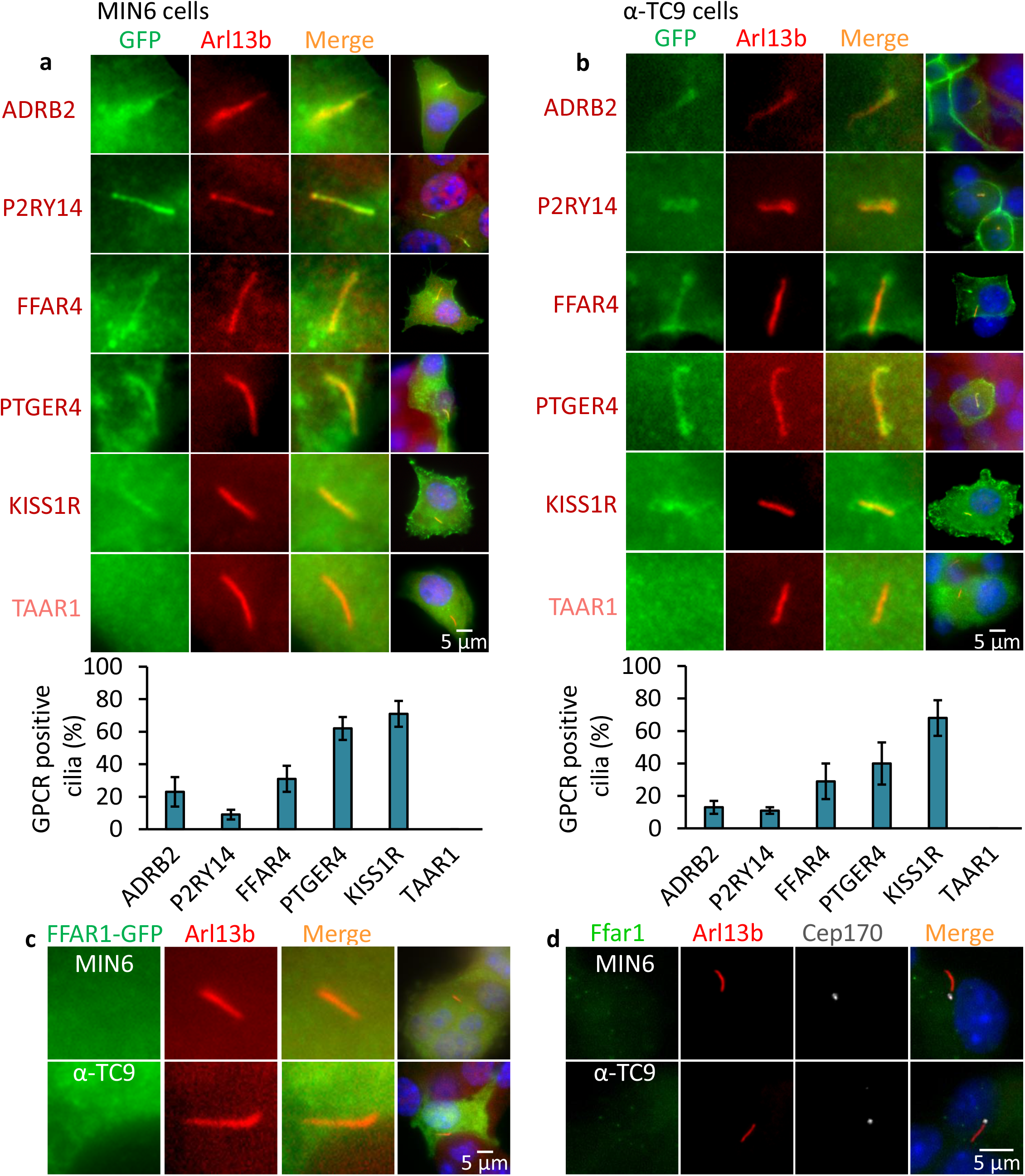
MIN6 (**a**) or α-TC9 (**b**) cells expressing GFP-tagged GPCRs were grown to confluence and immunostained with indicated antibodies. Percentages of GFP-positive ciliated cells (labelled Ar113b) are shown in (**a, b**) (down panel). Error bars in **a** and **b** represent mean ± s.d. (n = 3 independent experiments with 100 cells scored per experiment). (**c,d**) GFP-tagged FFAR1 (**c**) and endogenous FFAR1 (**d**) does not localize to the primary cilium of MIN6 and α-TC9 cells.

**Extended Data Fig. 2.**
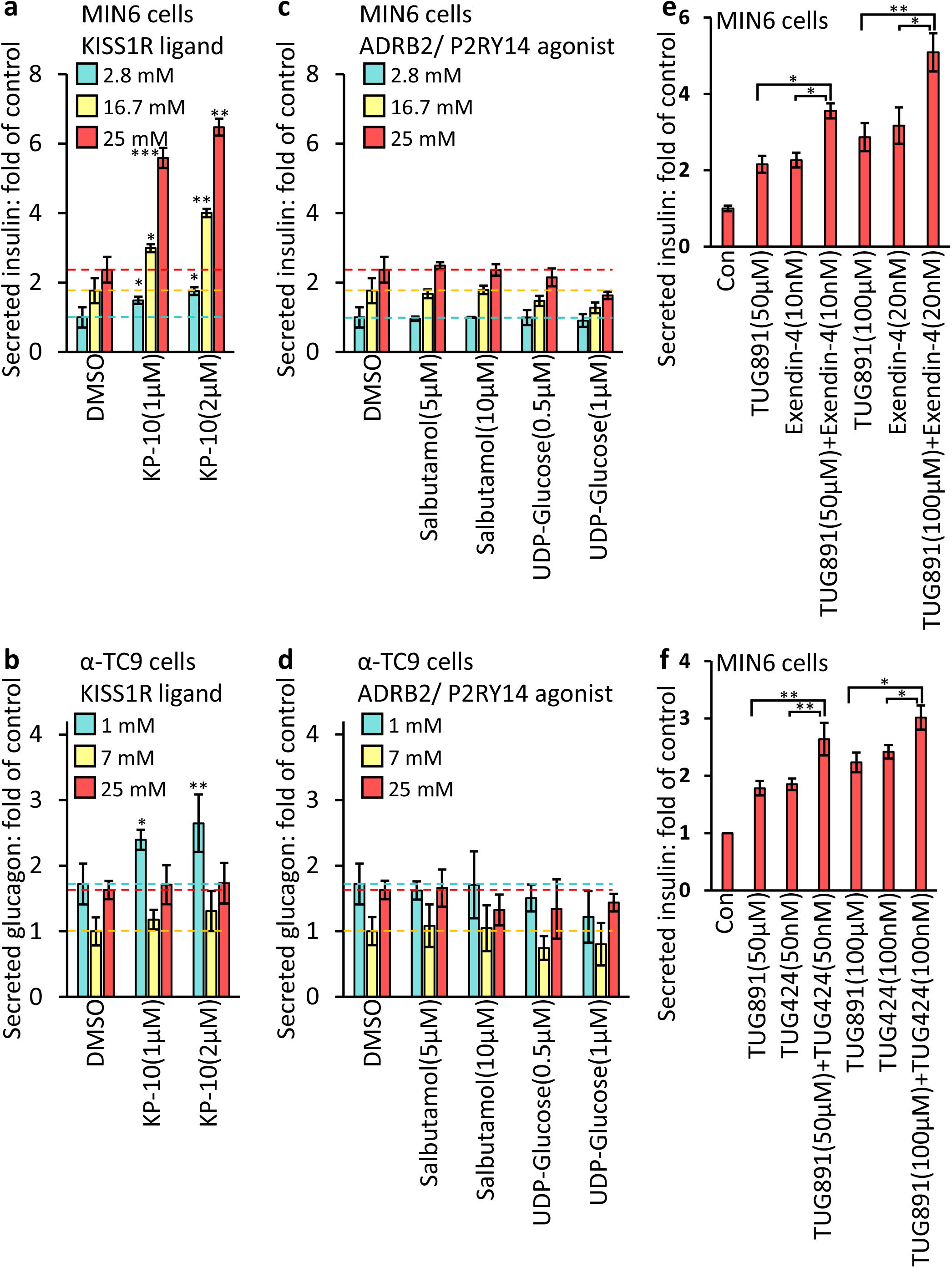
GSIS (**a,c,e,f**) and GSGS (**b,d**) induced by elevation (from 2.8 mM to 25 mM) or decrease (from 25 mM to 1 mM) of glucose levels and then effects of agonists on insulin or glucagon secretion have been evaluated. Bar graphs are normalized mean ± SD (n = 3 independent experiments); *p < 0.05; **p < 0.01; ***p < 0.001.

**Extended Data Fig. 3.**
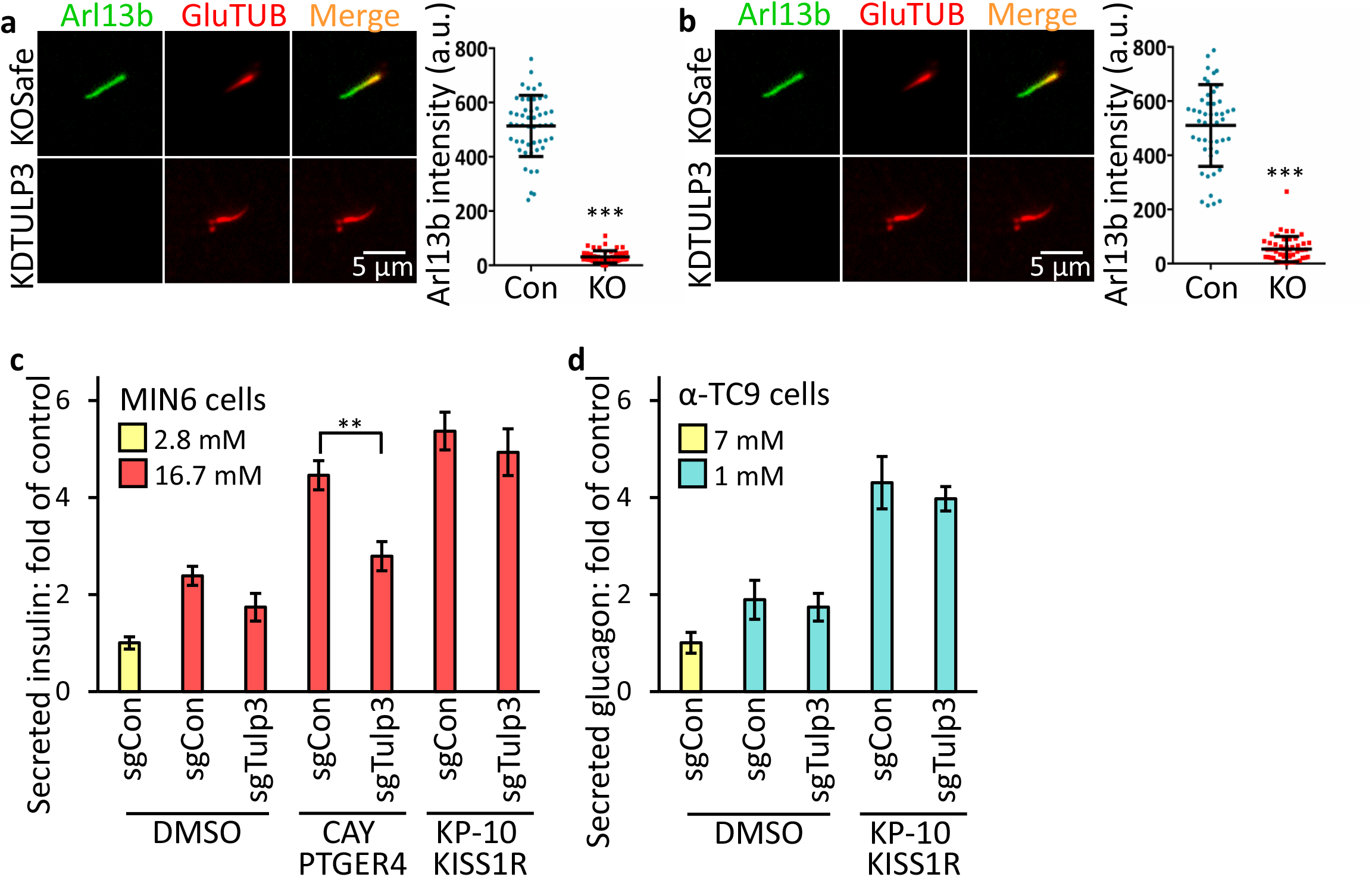
Loss of Tulp3 prevents ciliary Arl13b trafficking in MIN6 and α-TC9 cells. Control MIN6 (**a**) and α-TC9 (**b**) cells and Tulp3 knockout cell lines grown to confluence were immunostained with indicated antibodies. (**c**) Ptger4-regulated GSIS is cilia dependent. (**c,d**) Kiss1r-regulated GSIS (**c**) and GSGS (**d**) are cilia independent. Error bars in **c** and **d** represent mean ± s.d. (n = 3 independent experiments). *p < 0.05; **p < 0.01; ***p < 0.001.

**Extended Data Fig. 4.**
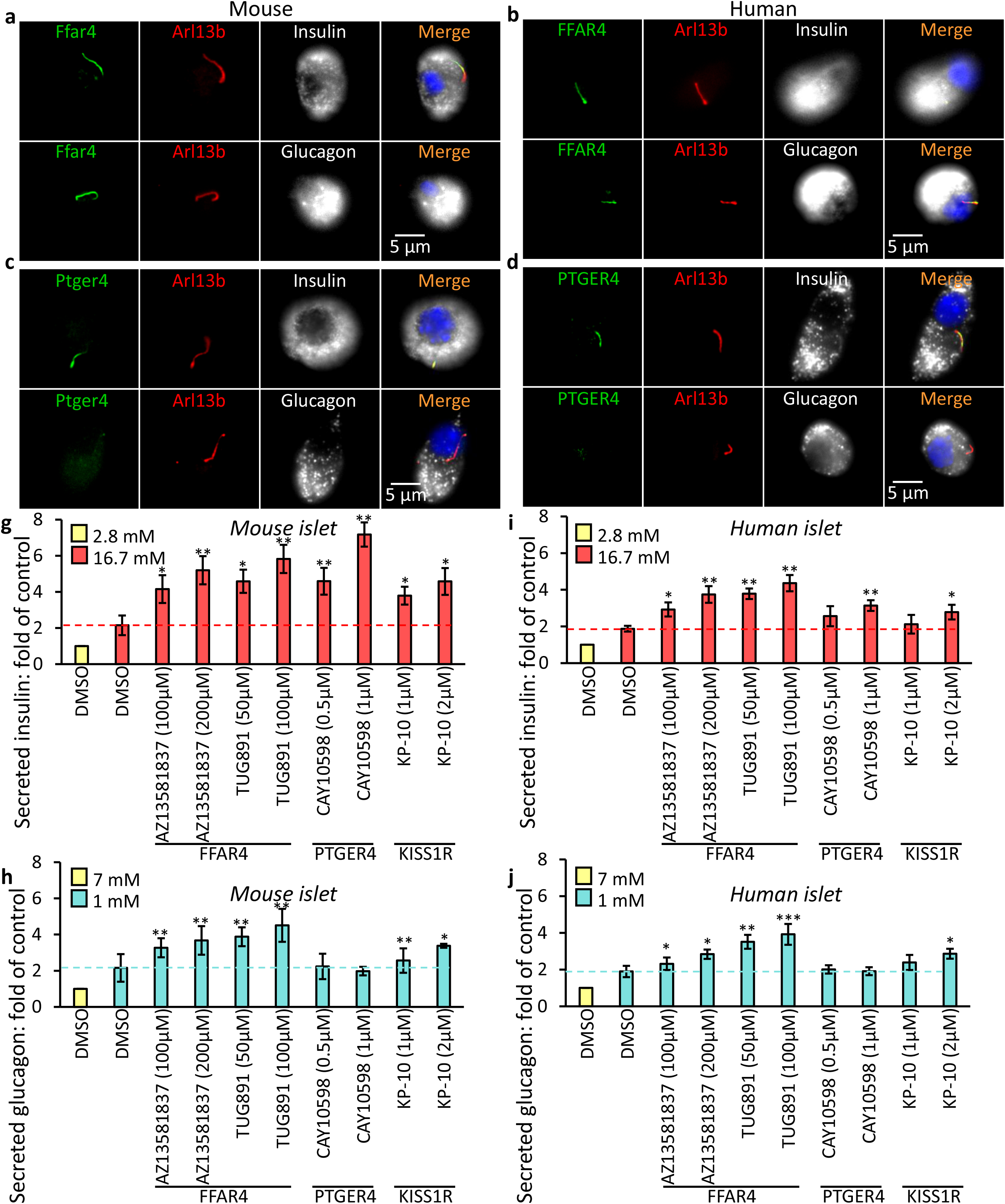
(**a-d**) Endogenous GPCRs localize to the primary cilium of dissected mouse (**a, c**) and human (**b, d**) pancreatic α- and β-cells. GSIS (**g, i**) and GSGS (**h, j**) induced by elevation (from 2.8 mM to 25 mM) or decline (from 25 mM to 1 mM) of glucose levels and then effects of agonists on insulin or glucagon secretion have been evaluated in mouse (**g, h**) and human (**i, j**) islets. Insulin and glucagon content of each treatment was measured using ELISA. Error bars in **g–j** represent mean ± s.d. (n = 4 independent experiments). *p < 0.05; **p < 0.01; ***p < 0.001.

**Extended Data Fig. 5.**
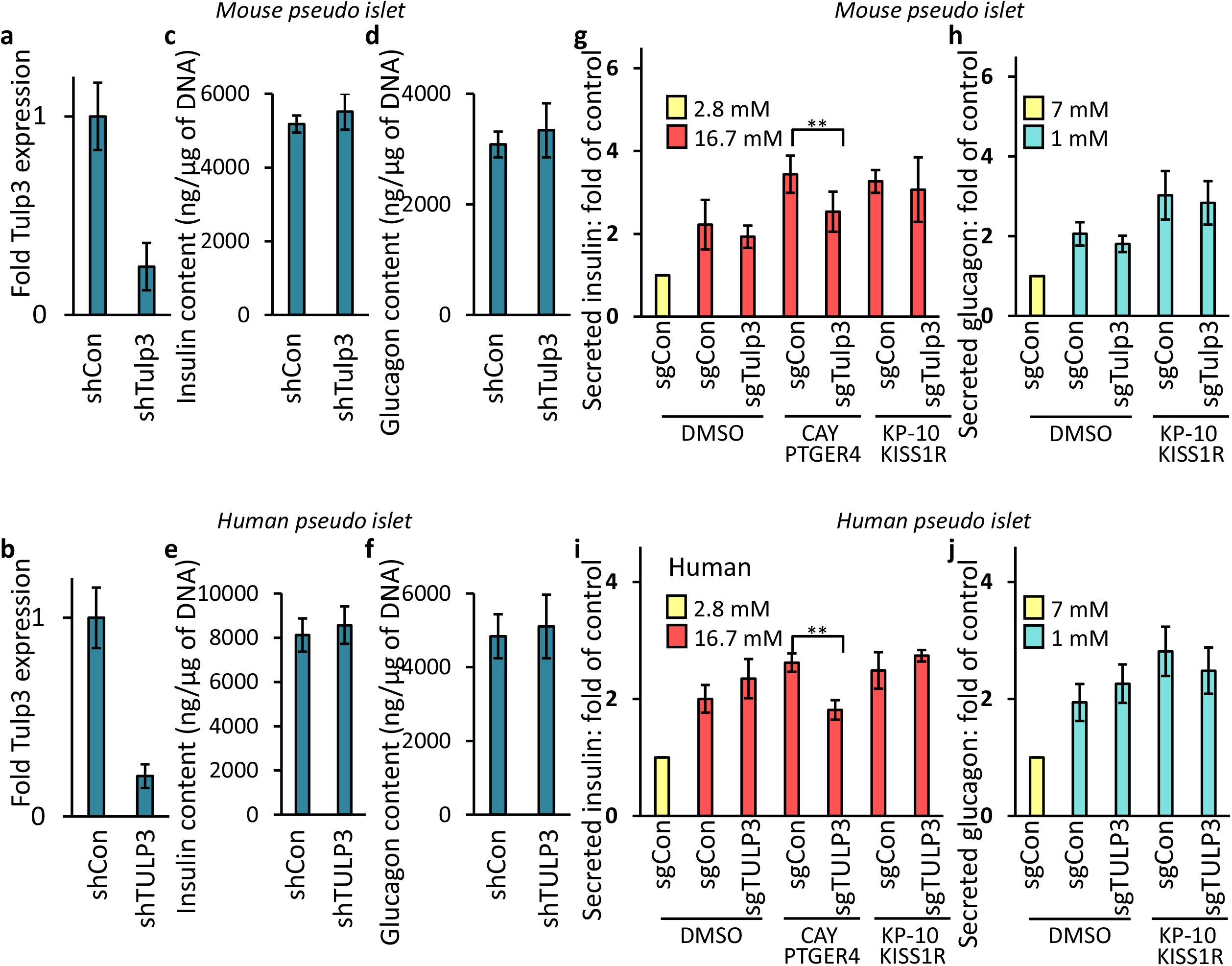
(**a, b**) Tulp3 mRNA expression in mouse (**a**) and human (**b**) pseudoislets following transduction with control or Tulp3 knockdown lentiviral vectors. (**c-f**) Insulin (**c, e**) and glucagon (**d, f**) content was unchanged in Tulp3 knockdown versus control mouse (**c, d**) or human (**e, f**) pseudoislets. (**g-j**) Ptger4-regulated GSIS is cilia dependent but Kiss1r-regulated GSIS and GSGS are cilia independent in mouse (**g, h**) and human (**i, j**) pseudoislets. Error bars in **a–j** represent mean ± s.d. (n = 4 independent experiments). *p < 0.05; **p < 0.01; ***p < 0.001.

**Extended Data Fig. 6.**
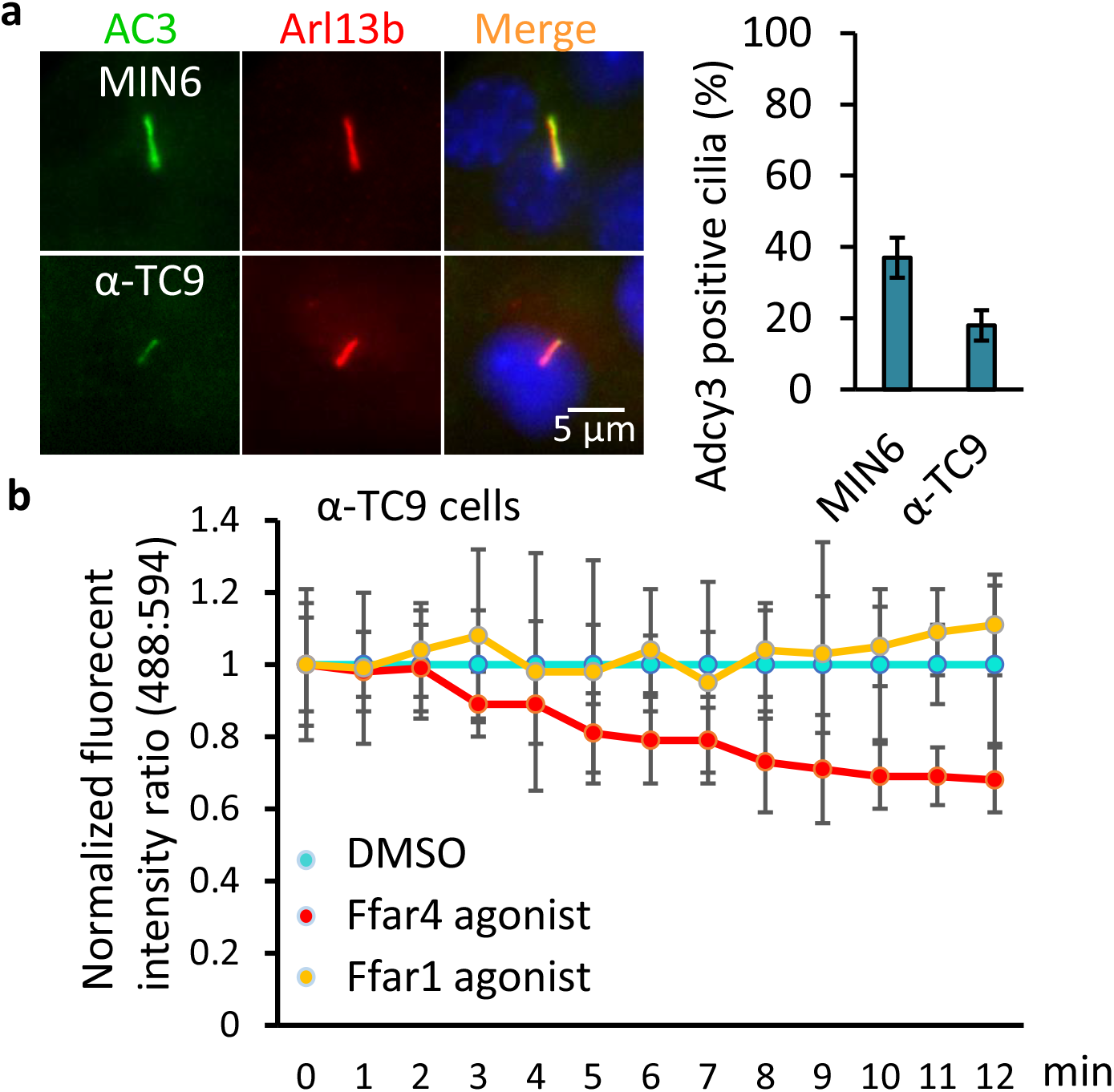
(**a**) ADCY3 (AC3) localizes to the cilia of MIN6 and α-TC9 cells. MIN6 and α-TC9 cells grown to confluence were immunostained with indicated antibodies. Percentages of AC3-positive ciliated cells (labelled Ar113b) are shown in (**a**, right). (**b**) Ffar4 regulates GSGS via cAMP in α-TC9 cells. Background subtracted ratio of fluorescence intensities are normalized to DMSO control and 0 s time point. Error bars in **a** represent mean ± s.d. (n = 3 independent experiments with 100 cells scored per experiment). *p < 0.05; **p < 0.01; ***p < 0.001.

## Notes

### Competing Interest Statement

The authors have declared no competing interest.

